# Hyperpolarization-activated cation channels shape the spiking frequency preference of human cortical layer 5 pyramidal neurons

**DOI:** 10.1101/2023.02.13.528352

**Authors:** Happy Inibhunu, Homeira Moradi Chameh, Frances K Skinner, Scott Rich, Taufik A Valiante

## Abstract

Discerning the contribution of specific ionic currents to complex neuronal dynamics is a difficult, but important, task. This challenge is exacerbated in the human setting, although the widely-characterized uniqueness of the human brain as compared to preclinical models necessitates the direct study of human neurons. Neuronal spiking frequency preference is of particular interest given its role in rhythm generation and signal transmission in cortical circuits. Here, we combine the frequency-dependent gain (FDG), a measure of spiking frequency preference, and novel *in silico* analyses to dissect the contributions of individual ionic currents to key FDG features of human L5 neurons. We confirm that a contemporary model of such a neuron, primarily constrained to capture subthreshold activity driven by the hyperpolarization-activated cyclic nucleotide gated (h-) current, replicates key features of the *in vitro* FDG both with and without h-current activity. With the model confirmed as a viable approximation of the biophysical features of interest, we applied new analysis techniques to quantify the activity of each modeled ionic current in the moments prior to spiking, revealing unique dynamics of the h-current. These findings motivated patch-clamp recordings in analogous rodent neurons to characterize their FDG, which confirmed that a biophysically-detailed model of these neurons captures key inter-species differences in the FDG. These differences are correlated with distinct contributions of the h-current to neuronal activity. Together, this interdisciplinary and multi-species study provides new insights directly relating the dynamics of the h-current to neuronal spiking frequency preference in human L5 neurons.

**Significance Statement:** Understanding the contributions of individual ionic currents to neuronal activity is vital, considering the established role of ion channel modifications in neuropsychiatric conditions. We combine *in vitro* characterization of the spiking frequency preference of human L5 cortical pyramidal neurons via the frequency-dependent gain (FDG) with new analyses of a biophysically-detailed computational model of such a neuron to delineate the connection between the dynamics of the hyperpolarization-activated cyclic nucleotide gated (h-) current prior to spiking and key properties of the FDG. By further determining that both these FDG properties and h-current dynamics are distinct in analogous rodent neurons, we provide convincing evidence for the key role of the h-current in the frequency preference of human L5 cortical neurons.

## Introduction

Discerning how ion channels contribute to neuronal output is a daunting but essential task, as specific channel types play key roles in neuropsychiatric conditions (Chang et al., 2019; Lupien-Meilleur et al., 2021; Martinez-Losa et al., 2018; Sun et al., 2019). This is particularly relevant to the study of the human brain considering its differences from preclinical models (Mohan et al., 2015; Eyal et al., 2016; Beaulieu-Laroche et al., 2018; Kalmbach et al., 2018; Boldog et al., 2018; Deitcher et al., 2017; Molnár et al., 2008; Verhoog et al., 2013; Kalmbach et al., 2021). Such differences motivate ongoing characterizations of human cell types (Moradi Chameh et al., 2021; Bakken et al., 2021; Kalmbach et al., 2021), a challenging task given limitations on human data (Rich et al., 2022) and the relative paucity of cell and circuit mapping tools used in rodent studies. Exacerbating this challenge is that ion channel activity is typically characterized using voltage clamp, preventing the simultaneous quantification of neuronal output. While an ion channel’s role can be inferred by comparing activity with and without channel blockade (Kostyuk et al., 1981; Ascoli et al., 2010; Thomas et al., 2007; Berger et al., 2001; Yue and Huguenard, 2001; Gu et al., 2005; Berger et al., 2003; Moradi Chameh et al., 2021; Kalmbach et al., 2018), these interpretations depend on the mechanism of blockade and potential compensatory interactions.

These considerations are especially pertinent when studying the brain’s oscillatory dynamics (Buzsaki, 2006). Understanding these rhythms’ genesis involves accounting for their specific frequencies, recording and analysis methods, disease, and functional correlates (Adamantidis et al., 2019; Buzsáki and Draguhn, 2004; Buzsáki et al., 2012; Hanslmayr et al., 2019; Maris et al., 2016; Mably and Colgin, 2018). Results from preclinical models, while invaluable, must be applied cautiously to the human brain (Mohan et al., 2015; Eyal et al., 2016; Beaulieu-Laroche et al., 2018; Kalmbach et al., 2018; Boldog et al., 2018; Deitcher et al., 2017; Molnár et al., 2008; Verhoog et al., 2013; Kalmbach et al., 2021); indeed, studies from human cortical slices have identified important nuances in their oscillatory dynamics (Florez et al., 2015; McGinn and Valiante, 2014).

Spiking frequency preference is an important factor underlying rhythm generation and signal transmission in cortical circuits (Higgs and Spain, 2009), being indicative of a cell’s potential role in pacemaking, phase-locking, and amplifying frequency-modulated inputs. Therefore, understanding the influence of ion channels on spiking frequency preference represents a significant contribution to understanding functionally important aspects of neuronal dynamics. Recent experimental work identified the frequency preference of human cortical layer 5 (L5) neurons via the frequency-dependent gain (FDG) (Higgs and Spain, 2009). These neurons exhibit a primary FDG peak between 2 and 6 Hz and a secondary peak at approximately double that frequency, both of which dissipate following application of the h-channel blocker ZD-7288 (Moradi Chameh et al., 2021), comporting with the hypothesis that the h-current contributes to low-frequency oscillatory activity (Beaulieu-Laroche et al., 2018; Kalmbach et al., 2018; Hu et al., 2002; Das and Narayanan, 2017). However, what specific features of h-current activity underlie its relationship with these oscillations remains mysterious.

Here, we dissect this complex neuronal dynamic in human cortical pyramidal cells by applying the FDG to computational models. We first show that a biophysically-detailed model of a human L5 neuron (Rich et al., 2021) reproduces key features of the experimental FDG, despite these features not being a constraint in model generation. Instead, this model was primarily constrained by subthreshold, h-current driven activity (Rich et al., 2021), meaning its reproduction of complex spiking activity quantified by the FDG represents a strong connection between the h-current’s dynamics and spiking frequency preference. Further supporting this conclusion is the similar response of the computational and experimental FDG to h-current blockade. By combining the Currentscape visualization tool (Alonso and Marder, 2019) with spike-triggered average (STA) analysis (Schwartz et al., 2006), we reinforce this connection by identifying distinctive properties of h-current activity contributing to FDG features.

We then carried out a similar exploration with rodent neurons, motivated by known differences between human and rodent h-currents (Rich et al., 2021; Moradi Chameh et al., 2021). If the h-current is of special importance to spiking frequency preference, we would expect a distinct FDG profile in rodent neurons; indeed, new patch-clamp experiments in rodent L5 cortical pyramidal cells identified a FDG profile distinct from analogous human neurons. We confirmed that a model rodent L5 pyramidal neuron (Hay et al., 2011) echoes these differences, justifying further application of our Currentscape/STA analysis. This revealed differences in the h-current’s contribution prior to spiking associated with the rodent model’s distinct FDG; considering the differing kinetics of the h-current in the human (Rich et al., 2021) and rodent (Hay et al., 2011; Kole et al., 2006) models, this indicates that the distinct dynamics of human h-channels are necessary for the frequency preference of human L5 neurons.

In summary, this study combines experimental results with novel *in silico* analyses to connect unique dynamics of the h-current to the spiking frequency preference of human L5 cortical pyramidal neurons. This interdisciplinary, multi-species approach provides convincing evidence for the h-current’s special effect on the spiking frequency preference of these neurons.

## Materials and Methods

### Computational models of human and rodent neurons

This work analyzes two computational models of cortical layer 5 (L5) pyramidal neurons, one of a human neuron (Rich et al., 2021) and another of a rodent neuron (Hay et al., 2011). Both models are accessible via ModelDB at senselab.med.yale.edu/ModelDB (Accession:266984 for Rich et al. (2021), Accession:139653 for Hay et al. (2011)). Our implementation of the rodent neuron model utilizes the parameters found in the *L5bPCbiophys3.hoc* file. Both models are implemented in the NEURON simulation environment (Carnevale and Hines, 2006) and contain a biophysically-detailed morphology, including a soma and various apical/basilar dendritic compartments.

Both models contain 10 ionic currents with differing maximum conductances, as well as different passive properties. The only differences in the kinetics of the ion channels are in the h-channel, the focus of the work of Rich et al. (2021). For ease of reference we refer to these currents by how they are denoted in the code of these models: *Na_Ta_t* is a fast, inactivating sodium current; *Nap_Et2* is a persistent sodium current; *K_Pst* is a slow, inactivating potassium current; *SKv3_1* is a fast, non-inactivating potassium current; *SK_E2* is a small-conductance calcium activated potassium current; *K_Tst* is a fast, inactivating potassium current; *Ca_LVA* is a low voltage activated calcium current; *Ca_HVA* is a high voltage activated calcium current; *Ih* is a non-specific hyperpolarization-activated cation current; and *Im* is a muscarinic current. We also track the passive current, denoted *pas*.

All *in silico* experiments are performed in the somatic compartments of the corresponding models.

### *In vitro* recordings

Data from human cortical L5 pyramidal neurons was obtained as described in Moradi Chameh et al. (2021). The data making up Figure 1**A-B** is taken from the associated open source dataset (Howard et al., 2022) and re-processed for presentation in a raw and normalized (see below) fashion. We also apply spike-triggered averaged analysis to the *n* = 3 human cells used to create Figure 5**f** in Moradi Chameh et al. (2021) to qualitatively assess the correspondence between this measure in the *in vitro* and *in silico* settings, specifically the comparisons before and after the application of ZD7288; this analysis had not been previously performed.

**Figure 1:**
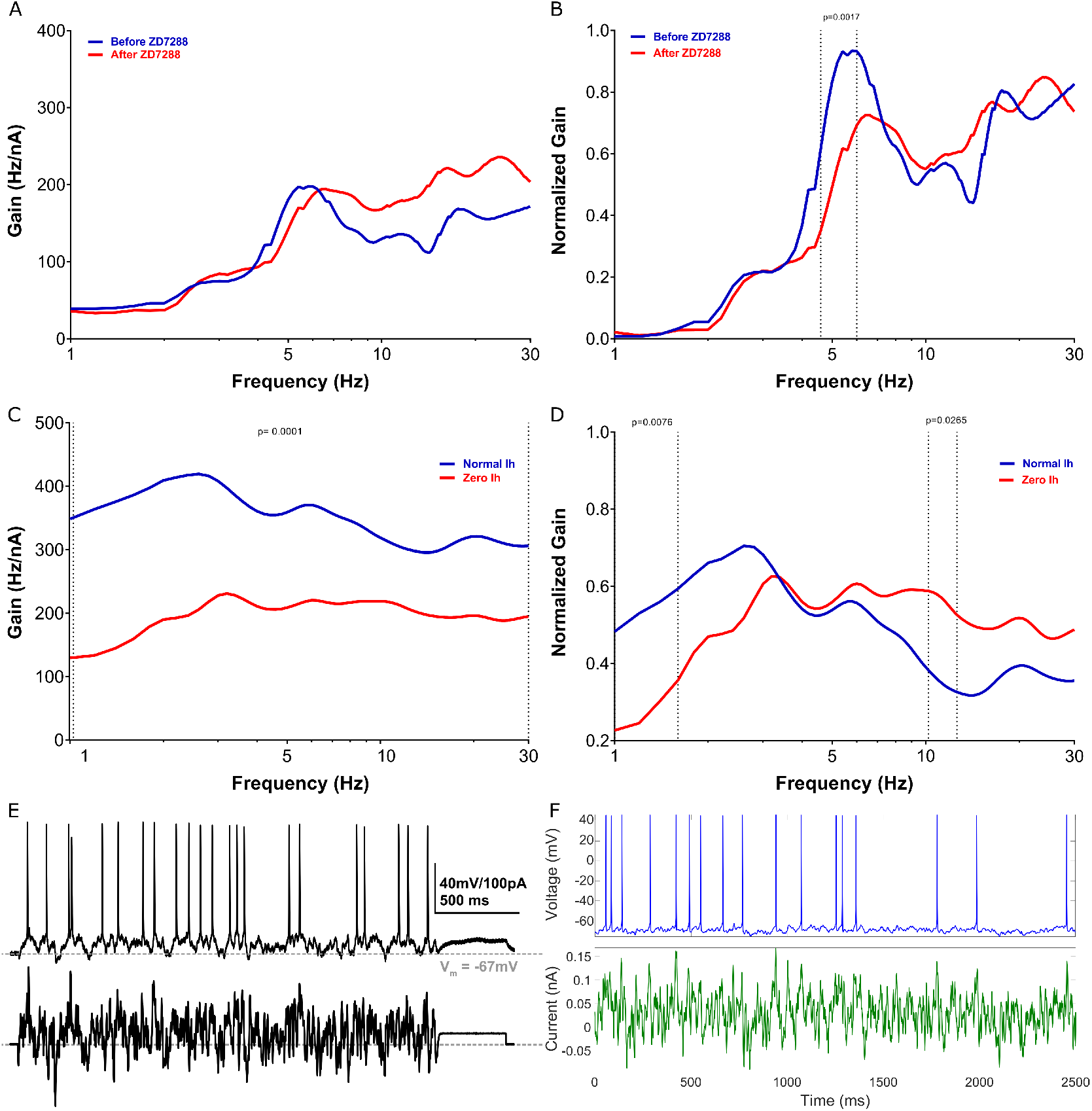
Model matches h-current mediated FDG peaks observed experimentally in human layer 5 cortical pyramidal neurons. **A-B** Experimentally calculated FDG (**A**) and normalized FDG (**B**) of L5 human pyramidal neurons with and without blockade of the h-channel with ZD-7288. Under control conditions, there is a clear low-frequency peak at approximately 5 Hz. After treatment with ZD-7288, the peak FDG values occur at >10 Hz. Normalized curves exhibit significant differences (p=0.0017) between 4.6 and 6 Hz, the location of this low-frequency peak. **C-D** FDG (**C**) and normalized FDG (**D**) derived from the model human L5 cortical pyramidal neuron under normal conditions and without h-current activity. Normalized plots emphasize the qualitative correspondence between the model and experimental settings: the model exhibits a low frequency peak (here at approximately 3 Hz) under normal conditions, while this peak dissipates (yielding a “flat” FDG profile for >3 Hz) without h-current activity. Non-normalized plots are significantly different for 1 to 30 Hz with p=0.0001; normalized plots are significantly different with p=0.0076 for 1 to 1.6 Hz and with p=0.0265 from 10 to 12.6 Hz. All significance values are derived from the 2-way ANOVA-Bonferroni’s multiple comparisons test. Note that the normalized plots (panels B and D) are *not* simply a rescaling of the absolute FDGs (panels A and C), but are processed as defined in the Methods. **E-F** Example input current (bottom) and output voltage (top) traces in the experimental setting (panel E, replicated with permission from Moradi Chameh et al. (2021)) and in the default model setting (panel F).

All experimental procedures involving mice were reviewed and approved by the animal care committees of the University Health Network in accordance with the guidelines of the Canadian Council on Animal Care. Mixed male and female wild type C57Bl/6J, age postnatal 21 days old were used for experiments. Mice were kept on a 12-hour light/dark cycle and had free access to food and water.

Brain slice preparation was done similarly for rodent tissue as outlined for human tissue in Moradi Chameh et al. (2021). Mice were deeply anesthetized by isoflurane 1.5-3.0%. After decapitation, the brain was submerged in (~4°C) cutting solution that was continuously bubbled with 95% O2-5% CO2 containing (in mM) sucrose 248, KCl 2, MgSO4.7H2O 3, CaCl2.2H2O 1, NaHCO3 26, NaH2PO4.H2O 1.25, D-glucose 10. The osmolarity was adjusted to 300-305 mOsm. Mouse somatosensory cortical slices (350 *μ*m) were prepared in the coronal plane using a vibratome (Leica 1200 V) in cutting solution. After slicing, slices were transferred to an incubation chamber filled with 32°C NaCl 126, KCl 2.5, NaH2PO4.H2O 1.25, NaHCO3 26, Glucose 12.6, CaCl2.2H2O 2, and MgSO4.7H20 which continuously bubbled with 95% O2–5% CO2. After 30 min, the slices were transferred to room temperature. Following this incubation, the slices were maintained in standard aCSF at 22–23°C for at least 1 h, until they were individually transferred to a submerged recording chamber (Moradi Chameh et al., 2021).

For electrophysiological recordings, cortical slices were placed in a recording chamber mounted on a fixed-stage upright microscope (Olympus BX51WI upright microscope; Olympus Optical Co., NY, USA). Slices were continuously perfused with carbogenated (95% O2/5% CO2) aCSF containing of (in mM): NaCl 123, KCl 4, CaCl2.2H2O 1.5, MgSO4.7H2O 1.3, NaHCO3 26, NaH2PO4.H2O 1.2, and D-glucose 10, pH 7.40 at 32-34°C. Cortical neurons were visualized using an IR-CCD camera (IR-1000, MTI, USA) with a 40x water immersion objective. Patch pipettes (3-6 MΩ) were pulled from standard borosilicate glass pipettes (thin-wall borosilicate tubes with filaments, World Precision Instruments, Sarasota, FL, USA) using a vertical puller (PC-10, Narishige). For somatic recordings of electrophysiological properties, patch pipettes were filled with intracellular solution containing (in mM): K-gluconate 135; NaCl 10; HEPES 10; MgCl2 1; Na2ATP 2; GTP 0.3, pH adjusted with KOH to 7.4 (290–309 mOsm). Data were collected with excitatory (APV 50*μ*M, Sigma; CNQX 25*μ*M, Sigma) and inhibitory (Bicuculline 10*μ*M, Sigma; CGP-35348 10*μ*M, Sigma) synaptic activity blocked.

Electrical signals were measured with a Multiclamp 700A amplifier and the pClamp 10.6 data acquisition software (Axon instruments, Molecular Devices, USA). Subsequently, electrical signals were digitized at 20 kHz using a 1440A digitizer (Axon instruments, Molecular Devices, USA). The access resistance was monitored throughout the recording (typically between 8-20 MΩ), and neurons were discarded if the access resistance was >25 MΩ.

For *in vitro* characterization of the frequency-dependent gain (see details below), a 2.5 s duration current stimulus of frozen white noise convolved with a 3-ms square function (Galán et al., 2008) was injected to each neuron 30 times for human neurons and 10 times for rodent neurons (with a 20 s inter-trial interval). An additional DC input, chosen specifically for each cell and experimental condition (i.e., before/after application of ZD7288) was injected to elicit approximately 12-15 spikes in the 2500 ms experimental paradigm. The FDG of the neuron is the average of the FDGs calculated for each trial. Data from n=6 rodent cells are shown in Figure 6.

### *In silico* experimental protocol

*In silico* experiments were designed to mimic those performed *in vitro*. Thus, all *in silico* results are analyses of a current-clamp injection of frozen white noise input (generated using the *makeNoise.m* MATLAB file included in our code repository with *σ* = 0.04 and *τ* = 3) plus a tonic “DC shift” chosen to elicit approximately 12-15 spikes in the 2500 ms experimental paradigm. In the human model, these experiments were performed in four scenarios: with the normal *Ih* maximum conductance as defined in Rich et al. (2021) (maximum somatic conductance of 5.14e-05 *S/cm*^2^), with that conductance doubled, with that conductance halved, and with that conductance zeroed (modeling channel blockade). Spikes were detected using a voltage threshold of greater than or equal to −55 mV for spike-triggered average analysis.

The DC shift for the human model was determined independently for each scenario given the effect of the h-current on the model’s excitability. These values were 0.003 nA for the scenario with doubled *Ih* maximum conductance, 0.031 nA for the default scenario, 0.040 nA when the maximum conductance was halved, and 0.073 nA when the maximum conductance was zero. In the rodent model, the DC shift was 0.38 nA both for the default model and the model with no h-current activity. The notable difference between the DC shift needed in the human and rodent models is reflective of these models’ differing excitabilities under this experimental protocol. While we would not expect a direct correspondence between the DC shifts in the *in vitro* and *in silico* settings for multiple reasons, including the effects of cell-to-cell variability (Golowasch et al., 2002; Marder and Goaillard, 2006) and the differing model development techniques used in Rich et al. (2021) and Hay et al. (2011), the increased DC shift necessitated *in silico* with decreased *Ih* maximum conductance in the human model reflects a similar trend before and after application of ZD7288 in human neurons *in silico*. The *in silico* DC shifts are also of similar orders of magnitude as those delivered *in vitro* in each setting.

All code used in this work is openly available at https://github.com/FKSkinnerLab/HumanL5FDG.

### Frequency-dependent gain

This work presents an in-depth investigation of the frequency-dependent gain (FDG) profiles of cortical L5 pyramidal cells in the human and rodent setting both *in silico* and *in vitro*. The methodology follows that of Higgs and Spain (2009), adapted to the *in silico* and *in vitro* setting as described above. This measure is used to quantify the propensity of a neuron to fire in phase with an oscillatory input of small magnitude relative to the overall input to the cell, or in essence to “track” particular frequencies in oscillatory input. The physiological implications of this measure include viable explanations for the oscillatory frequency preference of cortical regions, as outlined in detail in our previous work (Moradi Chameh et al., 2021). We note that this measures a distinct element of a neuron’s frequency preference compared to other common measures, such as subthreshold resonance in response to a ZAP input (Hutcheon and Yarom, 2000; Rich et al., 2021; Moradi Chameh et al., 2021).

After detecting action potentials, a time varying firing rate *r*(*t*) is calculated to be

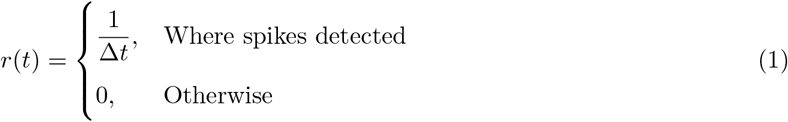

Then, the stimulus-response correlation (*c_sr_*) and the stimulus autocorrelation (*c_ss_*) is calculated

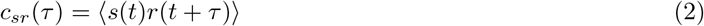

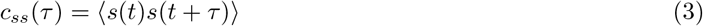

via for *τ* the time difference and *s*(*t*) representing the noisy stimulus.

The complex Fourier components *C_sr_*(*f*) and *C_ss_*(*f*) is then obtained, and the frequency-dependent gain is calculated as

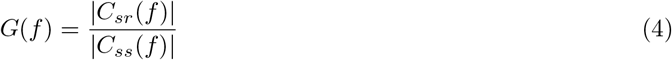

To derive averaged FDG plots for the model neurons, we generated 30 different 2500 ms frozen white noise inputs (as described above) to inject to the cell in addition to the tonic DC input. We then averaged the FDGs calculated from the outcomes of in silico current clamp experiments with each of these 30 different inputs. Note that this is distinct from the analogous *in vitro* protocol, in which the *same* noisy input is injected to each neuron multiple times (specifically, 10 trials for each of the 3 cells were done, and averaged for each cell and overall), motivated by the variable responses observed in single neurons subject to identical current injections. In contrast, the deterministic nature of both model neurons studied here would yield identical outputs in response to each injection of an identical current input. Considering these differences, we consciously chose our *in silico* technique to exploit the computational setting towards a more complete picture of the FDG of the model neurons in response multiple noisy inputs with the same statistics. While this choice might impair the exact correspondence between the *in silico* and *in vitro* settings, it does not affect the clear qualitative correspondence in primary features of the FDG that are the focus of our analyses.

Normalized FDG plots emphasize the location of the “peaks” that represent a frequency preference of these neurons, independent of the magnitude of the overall gain. A standard normalization 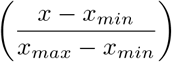 is performed over the 1-30 Hz range on the FDG resulting from each noisy input in the *in silico* setting and on the FDG resulting from each cell in the *in vitro* setting. These are then averaged to yield the normalized FDG for each setting and scenario.

The 2-way ANOVA-Bonferroni’s multiple comparisons test is used to derive statistical significance between FDG curves. A standard cutoff of p <0.05 is used to determine significance. Reported p-values are averaged over the regimes with significant differences.

### Currentscape visualization

In this work, we apply the Currentscape visualization tool (Alonso and Marder, 2019) to display the dynamics of ionic currents as percent contributions over time using stacked-area plots. This tool provides an interface between NEURON, where the neuronal cell model is situated and run, and Python, which facilitates data visualization. An array of currents within the model is inputted into Currentscape and split into inward and outward currents. Inward currents are represented as negative values while outward currents are represented as positive values. For each current, the percentage of its contribution to the overall inward/outward current at each time step is calculated.

During the simulation, the ionic currents, membrane voltage, and action potentials are extracted from the simulation in NEURON and imported into the Currentscape tool in Python. All code used in running these Currentscape visualizations are included in the code repository. Of note, whether the human (Rich et al., 2021) or rodent (Hay et al., 2011) model is implemented depends upon whether the command h.load file(“init_final.hoc”) or h.xopen(“Hay_setup.hoc”) is invoked. We set h.cvode active(0) to ensure a fixed time step for these simulations. The simulations performed in this study, motivated by the calculation of the FDG, involve calling the *noiserun* procedure with four inputs: the name of the file housing the noisy input, the value of *dt* (0.01 for human simulations, 0.33 for rodent simulations), the simulation duration (2500 ms), and the DC shift (outlined above for differing *Ih* conductances).

Ultimately, Currentscape outputs a plot containing: 1) Voltage traces (mV); 2) Total inward, outward, and total current traces (nA); and 3) percent contributions to the inward and outward current by each of the channels included in the model. Examples of these visualizations can be seen in Figures 2 and 3. Note that, in this study, the only compartment analyzed was the soma to best correspond with the *in vitro* experiments (Moradi Chameh et al., 2021).

**Figure 2:**
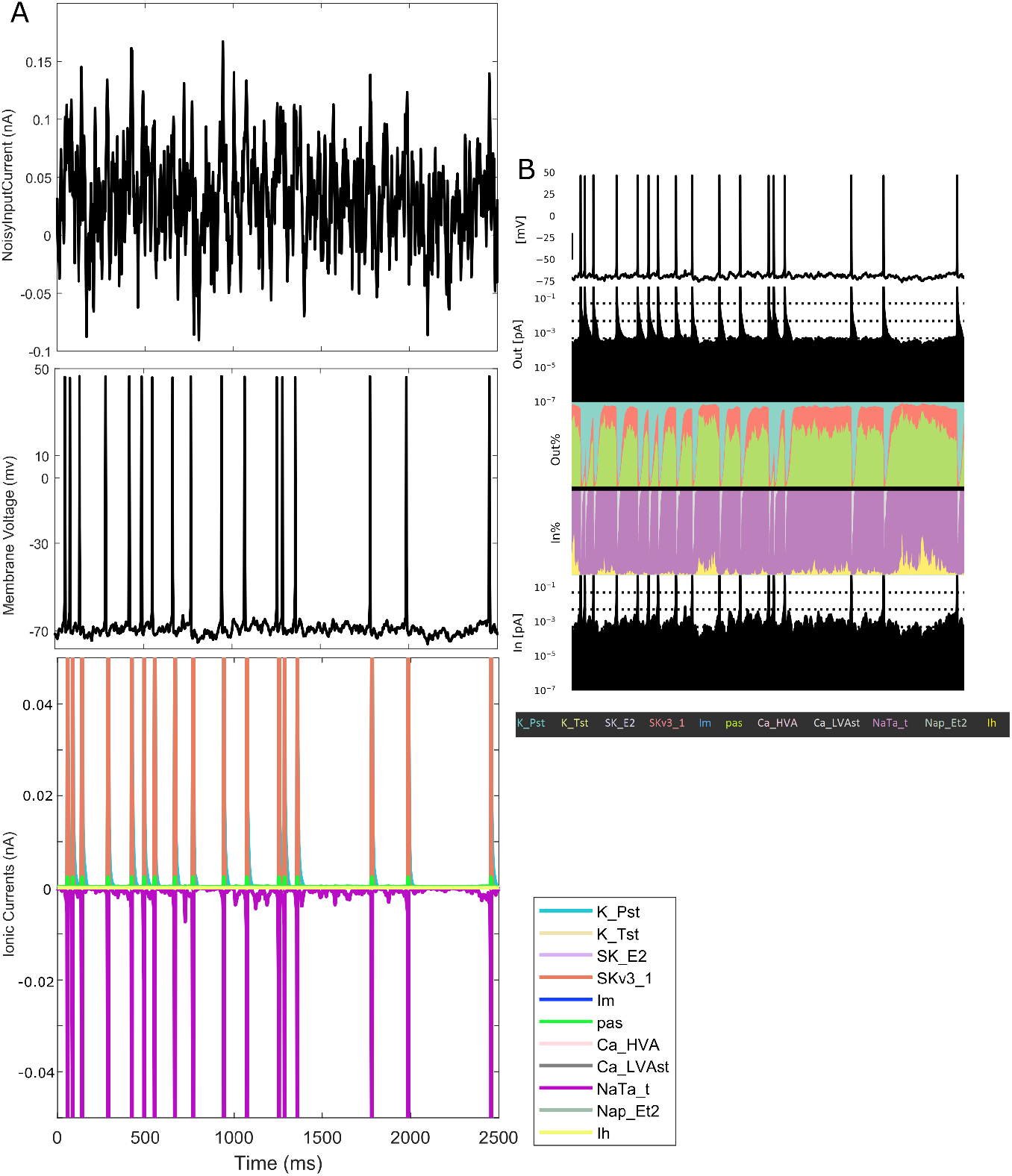
Currentscape visualization tool facilitates the identification of relative current contributions in spiking simulations. **A** Raw data obtained, via NEURON, of a simulation of a noisy input current injected into the L5 human pyramidal neuron model. Top to bottom: noisy input current, voltage trace and ionic current contributions. Of note is that the varying magnitudes of the ionic currents obscure one’s ability to jointly visualize their contribution to the neuron’s dynamics. **B** Using the Currentscape tool, the ionic current contributions are recontextualized as outward and inward percentage of contribution plots, allowing for a more intuitive visualization on a single scale despite the neuron’s spiking activity. Note that *Im* is not discernible in this visualization, denoting its minimal contribution to the dynamics of the model in this scenario.

**Figure 3:**
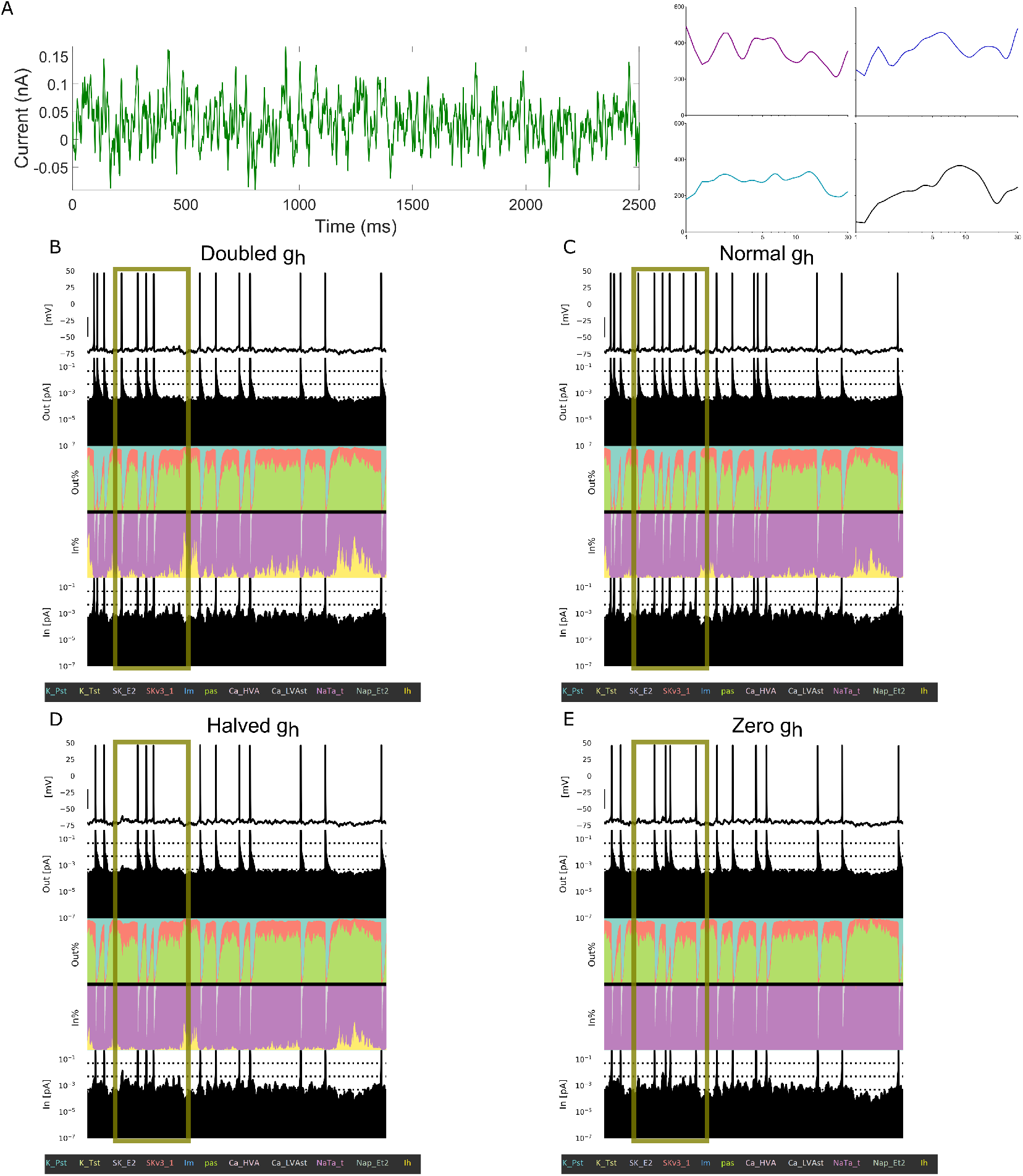
Currentscape visualization highlights differences in spiking modulated by h-current activity. **A** Left: The example noisy input current injected into the L5 human pyramidal neuron model with differing *Ih* conductance levels. Right: example FDGs (plot is Gain (nA) versus log-scaled Frequency (Hz)) from this input in each of four scenarios: with the *Ih* maximum conductance doubled (top-left, purple), normal (top-right, blue), halved (bottom-left, teal) or zero (bottom-right, black). **B-E** Currentscape visualizations corresponding with each of the scenarios outlined above. The h-channel contribution is indicated in the inward current contribution panel of the visualization in yellow, alongside the contributions of all the ionic currents and passive currents in the model. Spiking activity differs in each scenario (see highlighted regime) driven by the varying *Ih* conductance and the differing contributions of *Ih* to the neuron’s activity highlighted by this Currentscape visualization.

Currentscape simulations on the human neuron model were performed utilizing the Neuroscience Gateway (NSG), an online supercomputer (Sivagnanam et al., 2013). Identical code is used as outlined above and included in our code repository for simulations on NSG, but it must be packaged in one zip file included in the data section of the supercomputer. Once uploaded, performing the simulations on NSG involves executing a Python file that includes Currentscape and NEURON, as indicated above, in a specific platform of NSG that has NEURON via a Python interface.

Due to compatibility issues, Currentscape simulations on the rodent neuron model could not be performed on NSG. Running these simulations on personal machines required a coarser temporal mesh due to computational limitations: *dt* = 0.33 ms as opposed to *dt* = 0.01 ms used with the human neuron. This coarser discretization is responsible for the qualitative “jaggedness” of the rodent model’s FDG presented in Figure **6C-D**, but does not confound the features of that curve (the locations of FDG peaks) nor the visualization and analysis of the corresponding Currentscape plots.

### Spike-triggered average analysis

Traditionally, spike-triggered average analysis (Schwartz et al., 2006) is used to quantify the average input current preceding spiking activity in an neuron, as was done in L5 human cortical pyramidal neurons by Moradi Chameh et al. (2021) using a 30 ms window. Here, we exploit the additional data afforded by Currentscape to perform analogous analysis on the percent contribution to the inward or outward current of individual channels preceding spiking in our neuron models. We analyze these dynamics both in the 30 ms window used by Moradi Chameh et al. (2021) and in longer 100 and 200 ms windows to ensure capture of the slow h-channel kinetics.

To perform these analyses, we first followed the *in silico* experimental protocol outlined above. We detected spikes and only considered those for which there was no other spike in the 30, 100, or 200 ms preceding it (as appropriate for the given time window). We averaged the dynamics of the percent contributions of the various currents over the time window prior to each of these spikes for each of the 30 noisy input currents. In this way, we extend the oft-used experimental tool of spike-triggered averaging beyond the input current, using Currentscape, to describe the dynamics of individual ionic and passive currents in the moments prior to spiking. This allows us to directly discern whether any ionic current contributes uniquely to spike timing.

#### Quantification of STA dynamics

We performed additional analysis to quantify the qualitatively different features of the STAs of Currentscape derived percent contributions to inward/outward currents. These STAs are presented in Figures 4-6; for a particular current *c*, denote the raw STA trace as 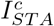. We process 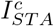 by first normalizing it over the given time window, dividing each value by the maximum value for outward currents or minimal value for inward currents in said window. We then “smooth” the curve using MATLAB’s built in *smooth* function (MATLAB, 2019), yielding a processed trace, denoted 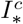, that ranges between 0 and 1.

**Figure 4:**
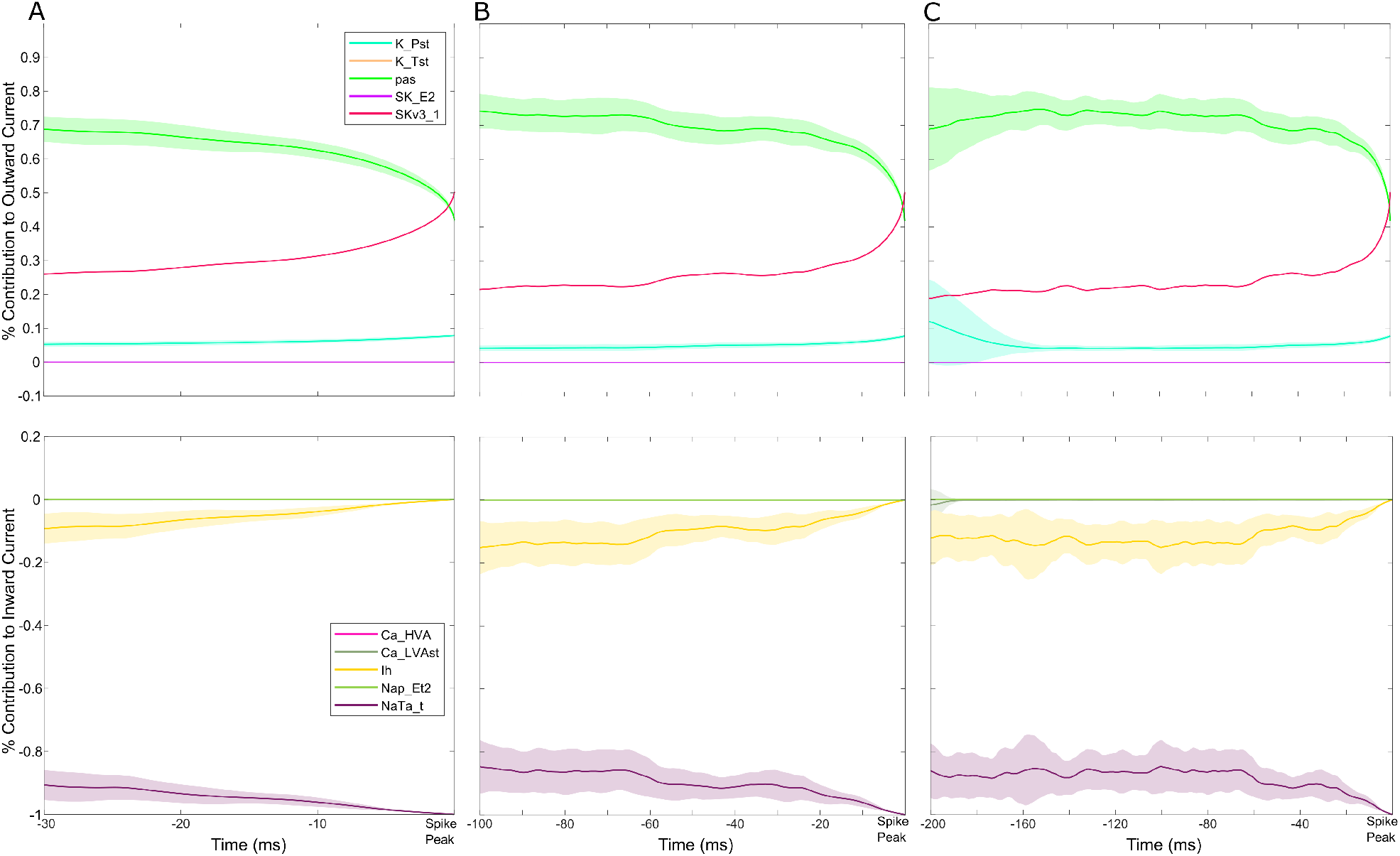
Currentscape data processing merged with spike-triggered average analysis of human cortical layer 5 neuron dynamics. **A-C** Representations of the STA for the percent contribution to outward (top panel) and inward (bottom panel) currents for 30 ms (panel C), 100 ms (panel D), and 200 ms (panel E) prior to a spike. A unique dynamic is observed in the dynamics of the h-current in panel E: over these 200 ms, the contribution of the h-channel in distinctively non-monotonic. The shaded portion of each plot represents ± one standard deviation over the 30 repetitions with distinct noisy inputs. *Im* is not included in these plots given its minimal contribution to model dynamics (see Figure 2).

How 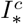 changes approaching spiking reflects its relative contribution and subsequent control of the spiking process. In particular, we characterize the monotonicity of 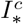. Mathematically, a monotonic function is one that either never increases or never decreases. Here, we quantify the monotonicity of 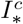 by calculating the proportion of the time series that is increasing and decreasing: a time series that is either increasing or decreasing 100% of the time is purely monotonic, while one that decreases 50% of the time and increases the other 50% would be maximally non-monotonic (i.e., it would be increasing and decreasing for equal proportions of the time window). To accomplish this, we calculate the difference quotient for each time step, yielding a new time series 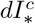. The proportion of negative values in this time series yields the proportion of the 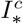 time-series that is decreasing, and vice versa, which we use to draw conclusions about the monotonicity of the current *c*’s contribution to the inward/outward current prior to spiking.

## Results

### Model replicates key features of experimentally derived FDG with and without h-current contribution

The frequency dependent gain (FDG) identifies the propensity of a neuron to spike in phase with a small oscillatory input relative to the neuron’s net input (Higgs and Spain, 2009). Experimentally, as shown in Figure 1**A-B**, human L5 cortical pyramidal neurons exhibit a primary FDG peak in a low frequency range between 2 and 6 Hz, with a secondary peak at approximately double that frequency. Under h-current blockade with ZD-7288, these peaks dissipate and the gain mostly increases with increasing frequency. This is quantified by the significant difference (p=0.0017) between 4.6 and 6 Hz in the normalized curves (achieved via processing the raw FDG data as outlined in the Methods), a range including the low-frequency peak. An example of the experimental traces used to derive the FDG is illustrated in Figure 1**E**.

Notably, the FDG profile of the human L5 model neuron presented by Rich et al. (2021), illustrated in Figure 1**C-D**, replicates key features of the experimental FDG: there is a low-frequency peak (here at around 3 Hz) and a secondary peak at double this frequency. These peaks also dissipate when the h-current is blocked in the model neuron, including a “flat” FDG profile for > 3 Hz. The raw FDG traces with and without h-current activity (panel C) are significantly different with p=0.0001 for the entire frequency range. Normalized plots (achieved via processing the raw FDG data as outlined in the Methods) emphasize the presence (or lack thereof) of peaks that indicate a frequency preference: these are significantly different for 1 to 1.6 Hz with p=0.0076 and for 10 to 12.6 Hz with p=0.0265. The difference at above 10 Hz is particularly notable as it shows that the decrease in gain at high frequencies in the default model, which serves to accentuate the peaks indicating frequency preference, is significantly diminished without h-current activity. An example of the model output under the paradigm used to calculate the FDG is illustrated in Figure 1**F**.

Considering the model neuron was constrained primarily by subthreshold voltage traces and not by spiking activity in response to a noisy input current (Rich et al., 2021), the correspondence between the primary qualitative features of the experimental and model FDG profiles is a non-trivial and important result. Given the focus of the modeling process in Rich et al. (2021) on capturing h-current activity, it stands to reason that this FDG correspondence is due in part to the model’s accurate encapsulation of h-current dynamics. This conclusion is supported by the additional correspondence between the FDG profiles of the experimental neuron treated with ZD-7288 and the model neuron with no h-current activity. While the similarities between the *in silico* and *in vitro* FDGs are not exact, this is to be expected given the model generation process described above, as well as the different processes for generating averaged FDGs (averaging over different cells in the *in vitro* setting versus averaging over different noisy inputs in the *in silico* setting; see details in Methods) necessitated by the deterministic nature of the model neuron. One discrepancy of note is the increased gain at high frequencies seen experimentally following the application of ZD-7288; while the decrease in gain at these frequencies is significantly diminished in the model without h-current activity, an overall increase in gain is not observed. This is likely a consequence of secondary effects of treatment with ZD-7288 (Poolos, 2004; Sánchez-Alonso et al., 2008; Chevaleyre and Castillo, 2002), including an increased input resistance, that are not encapsulated by simply removing h-current activity in the model neuron. Nonetheless, the correspondence between the primary FDG features in the control setting (low-frequency peak and secondary peak at approximately double the frequency) and with h-current blockade (suppressed low-frequency activity and lack of a notable decrease in gain at > 10 Hz) justifies the use of this *in silico* model as a tool to better understand the contribution of the h-current to the neuron’s FDG.

### Currentscape visualization of noisy current injections highlights current contributions

We sought to fully exploit the *in silico* setting to perform measurements that could not be done *in vitro*: simultaneously quantifying the activity of each ionic current alongside the voltage response to a noisy input current. Currentscape (Alonso and Marder, 2019) is invaluable for the visualization of these current contributions within spiking simulations. As illustrated in Figure 2**A**, current activity differs by orders of magnitude between neuronal spiking and subthreshold activity, obscuring the evolution of these contributions over the course of a simulation. Currentscape addresses this problem, as illustrated in Figure 2**B**, by instead calculating the percent contribution of a given current to the net inward/outward current at a particular time step. This provides a more intuitive visualization of the evolution of ion channel contributions on a single scale despite the spiking dynamics of the neuron.

We employed this tool to decipher and highlight differences in neuronal dynamics as the h-current’s contribution is changed (by altering its conductance value) in Figure 3. An example of the noisy input current used in these simulations is illustrated in Figure 3**A** (note the average value of this input, its “DC Shift”, is altered in each setting reflecting changes in the *Ih* maximum conductance; see details in Methods), and Currentscape plots are illustrated when the *Ih* maximum conductance is doubled (Figure 3**B**), at its default value (Figure 3**C**), halved (Figure 3**D**), and zeroed (Figure 3**E**). Currentscape visualization allows for dynamics of the h-current (yellow inward current) to be discerned that would otherwise be obscured by examining raw current magnitudes as in Figure 2**A**.

We highlight a regime in panels B-E with a gold box in which spiking differs in each scenario. Using Currentscape visualization, we can identify increased h-current activity at the end of this period when the *Ih* maximum conductance is doubled and when it is halved (panels B and D) compared to both the default state and when there is no h-current activity (panels C and E); in the former scenarios there is no spiking during this period, while in the latter scenarios there is. This non-linear relationship between h-current activity, h-current conductance, and spiking highlights the complexity of these interactions in this biophysically-detailed neuron model, and motivates further quantifications in search of clearer relationships.

There are multiple ways one might quantify the differences in these spike trains as a result of the varying contributions of the h-current. One is the FDG itself; we include FDG plots of these specific example trials in Figure 3 to illustrate how FDG properties differ for differing spike trains. Differences are apparent from the FDGs of these individual trials, and the differences between FDGs in these scenarios averaged over multiple trails are statistically examined below. Another commonly used quantification of spike train features is the coefficient of variation (CV) of the inter-spike intervals (ISIs); while this is commonly measured in response to tonic current steps experimentally (Angelo and Margrie, 2011), here we apply this measurement to the response of our model to noisy input. We calculated the CV of the ISIs in each of the four scenarios for all 30 trials, and found the CV of the ISIs to be 0.78 for double *Ih* maximum conductance, 0.81 for normal *Ih* maximum conductance, 0.92 for halved *Ih* maximum conductance, and 1.05 for zero *Ih* maximum conductance. These CVs are significantly different between the doubled and halved *Ih* maximum conductance scenario, the doubled and zeroed *Ih* maximum conductance scenario, and the normal and zeroed *Ih* maximum conductance scenario (p<0.05 in the first case, p<0.001 in the last two cases; two sample coefficient of variation test). This pattern follows from the role of the h-current in dictating a spiking frequency preference: a neuron with a pronounced FDG peak will better track one of the many sinusoidal frequencies making up the white noise input, in turn firing more regularly and with less variability (smaller CV). This is borne out by the experimental data as well: analysis of the data yielding Figure 1**A-B** shows that the CV of the ISIs is significantly increased following blockade of the h-current with ZD-7288 (0.413 vs. 0.333; p=0.00004, two-sample coefficient of variation test). The FDG and CV of the ISIs can thus be considered complimentary measures of a spiking frequency preference, representing further quantitative support for the notable differences in the spiking activity of the model neuron in response to varying *Ih* maximum conductance.

These relationships imply that the h-current affects more than the neuron’s basic excitability features (Dyhrfjeld-Johnsen et al., 2009; Gasselin et al., 2015; Hogan and Poroli, 2008): distinct spike trains, associated with distinct time-courses of h-current activity, are observed for varying *Ih* maximum conductances despite compensation for changes in excitability via the DC input. Indeed, the contribution of the h-current to neuronal spiking appears more complex, causing neurons with differing *Ih* maximum conductances to respond differently to the same noisy input, often in unintuitive ways (i.e., the lack of spiking towards the end of the highlighted region seen when the *Ih* maximum conductance is doubled *and* halved, compared to multiple spikes in the default setting).

Importantly, these conclusions would be difficult to directly establish using only contemporary *in vitro* experimental techniques, and likely obscured without Currentscape visualization in the *in silico* setting. They remain, however, entirely qualitative. In the following, we further exploit the “percent contribution” measure generated by Currentscape to quantitatively explore the relationship between h-current activity and spiking activity.

### STA analysis quantifies unique dynamics of the h-current before spiking

To quantify the contributions of each ionic current to spiking activity, we adapted spike triggered average (STA) analysis (Schwartz et al., 2006) by applying this technique to the percent contribution of individual ionic currents to the net inward and outward current derived using Currentscape. In this process, we also confirmed that the traditional STA, as applied to the noisy current input (as in Moradi Chameh et al. (2021)), is qualitatively similar in our model and in the human neurons studied in Figure 1**A-B** (results not shown). This represents an additional validation that the model reasonably captures important features of the spiking dynamics of human L5 cortical pyramidal neurons.

We applied STA analysis to each individual current’s percent contribution to the net inward/outward current at each time step, as quantified using Currentscape. We visualized this measure for 30 ms (Figure 4**A**), 100 ms (Figure 4**B**), and 200 ms (Figure 4**C**) windows prior to spiking, with the larger time windows motivated by the slow kinetics of the human h-current (Rich et al., 2021). Differences in the h-current’s dynamics relative to other currents became apparent in the 200 ms prior to spiking, during which we see that the h-current’s contribution to the inward current is non-monotonic: on average, the percent contribution increases in the 200 and 100 ms prior to spiking, but then decreases towards zero until the spike peak (this monotonicity is further emphasized by the “zoomed in” plot in Figure 5**B**). In contrast, each of the outward currents behaves approximately monotonically for a larger proportion of the 200 ms time frame and exhibits more variability in the regime in which it does not: the *pas* current largely decreases, the *K_Pst* and *SKv3_1* currents largely increase (the *SKv3_1* current with notably minimal variability), and the *K_Tst* and *SK_E2* currents both have a minimal, near zero percent contribution. Meanwhile, amongst the inward currents, *Ca_HVA*, *Ca_LVAst*, and *Nap_Et2* all have near-zero percent contribution in the 200 ms time frame. This leaves the percent contribution of the *NaTa_t* current inversely proportional to the percent contribution of the h-current, as these currents together account for ~ 100% of the inward current. Considering the h-current is maximally active at sub-threshold voltages and acts on a slower time scale, while the *NaTa_t* current contributes primarily to action potential generation at a very fast time scale (Rich et al., 2021), we can reasonably assume that most changes in the inward current prior to spiking are driven primarily by the h-current.

**Figure 5:**
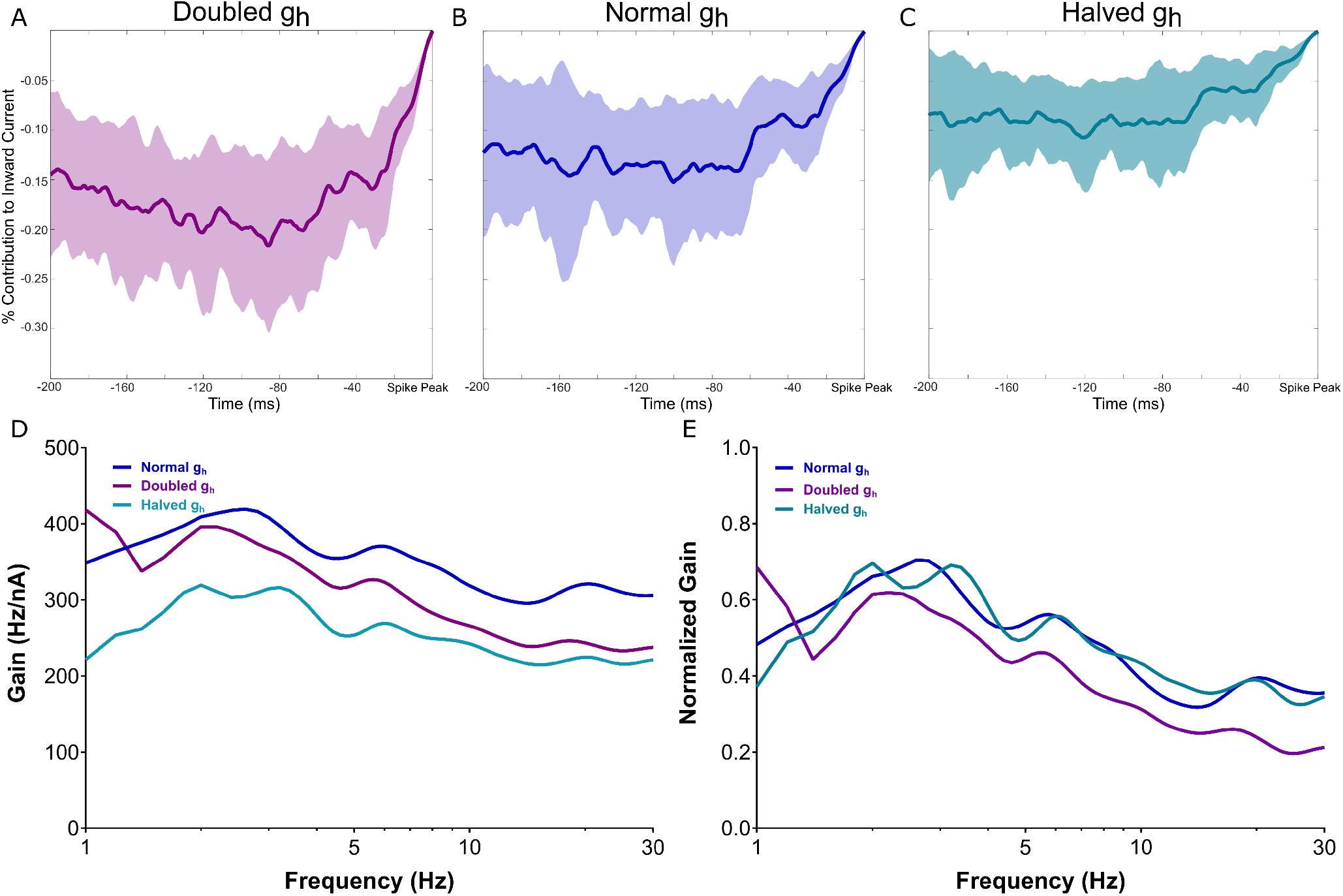
The non-monotonic nature of h-current activity prior to spiking is exaggerated with an increased h-channel conductance and diminished with a decreased h-channel conductance. **A-C** Visualizations of the h-channel contributions preceding spiking with the *Ih* maximum conductance doubled in panel A, normal in panel B, and halved in panel C, with the shaded regions representing ± one standard deviation. **D-E** Non-normalized (**D**) and normalized (**E**) FDGs in each of the above scenarios. The normalized plots highlight the “flatter” decay of the FDG at high frequencies when the *Ih* maximum conductance is halved and the more precipitous drop when the *Ih* maximum conductance is doubled. The former diminishes the influence of any low-frequency peaks, while the latter accentuates it.

Following the process outlined in the Methods, we quantified the non-monotonicity of the h-current’s contribution to the inward current visualized in the 200 ms STA. Given the relationship between the percent contribution of the *Ih* and *NaTa_t* currents described above, we focused on comparing the 200 ms STA of the h-current to the 200 ms STA of the outward ionic currents; indeed, the primary ionic currents contributing to the outward current, *SKv3_1* and *K_Pst*, both have quantitative distinctions from the dynamics of the h-current. The former current is almost entirely monotonic: 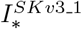 is increasing for 94.72% of the 200 ms prior to spiking, in sharp contrast with the h-current, for which 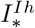 is decreasing for just 62.50% of the 200 ms prior to spiking. Meanwhile, there is a noticeable increase in the variability of the contribution of *K_Pst* between 200 and 150 ms prior to spiking (see the fainter ± STD shading), a clear distinction in comparison to the other ionic currents. The remaining ionic currents either contribute minimally in the 200 ms prior to spiking (*Ca_HVA*, *Ca_LVAst*, *Nap_Et2*, *K_Tst*, *SK_E2* ; also *Im* which is omitted from the plots as mentioned in Figure 3) or have a percent contribution approximately inversely proportional to that of the h-current (*NaTa_t*). Thus, the qualitatively observed uniqueness of the h-current’s 200 ms STA relative to other ionic currents is supported by quantitative analysis.

Next, we applied our STA analysis to three of the scenarios studied in Figure 3; double, normal, and halved *Ih* maximum conductance. As can be seen in Figure 5**A-C**, the curvature of the STA of the h-current’s percent contribution to the inward current is exaggerated when its conductance is doubled, and diminished when it is halved. We quantified this by examining 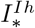 in each scenario; we found that 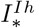 was decreasing for 70.37% of the STA time-series when the *Ih* maximum conductance was halved, 62.50% of the STA time-series for this conductance’s default value, and just 55.89% of the STA time-series when this conductance was doubled. This pattern confirms quantitatively what we observed qualitatively, that the monotonicity of these curves is affected by the amount of h-current activity, as dictated by the *Ih* channel’s maximum conductance. These changes, elicited solely by alterations to the h-current’s maximum conductance, are also confirmatory evidence that the features of the *NaTa_t* current’s percent contribution are most likely attributable to its relationship with the *Ih* current’s percent contribution, as discussed above.

These differences correspond with alterations to the corresponding FDGs, displayed non-normalized in Figure 5**D** and normalized in Figure 5**E**. Notably, when the *Ih* maximum conductance is halved, the low frequency peak appears “split,” which undermines a key feature of the *in vitro* and default *in silico* FDG: the presence of a single low-frequency peak and single secondary peak at approximately double the frequency. This can also be seen in the individual FDGs in Figure 3**A**. Additionally, the decay of the gain at high frequencies is notably more gradual, potentially mitigating the functional frequency preference quantified by the FDG. Meanwhile, when the *Ih* maximum conductance is doubled, the decay of the gain at high frequencies is more precipitous, strengthening the neuron’s frequency preference at the low-frequency peaks.

The qualitative differences described above are confirmed via quantitative statistical testing. The normalized FDG when the *Ih* maximum conductance is doubled is significantly different from the default curve, for p<0.05, for 19.2-30 Hz, while the non-normalized FDGs are significantly different for 6.4-30 Hz. Meanwhile, the non-normalized FDG when the *Ih* maximum conductance is halved is significantly different from the default curve, for p<0.05, for the entire 1-30 Hz range.

We note that the idiosyncratic maximum gain at 1 Hz when the *Ih* maximum conductance is doubled is a likely side effect of the model’s increased excitability in this scenario (as noted in the Methods, the DC shift necessary to elicit appropriate spiking is smaller by an order of magnitude when the *Ih* maximum conductance is doubled as compared to the default model). While this might be compensated for by changing other elements of the model neuron, we consciously chose to *only* manipulate the h-current in order to minimize any potential confounds. This nuance emphasizes the need to appropriately contextualize the conclusions drawn when altering model neurons such as these (see the discussion of “hybrid” model neurons in Rich et al. (2021)).

This analysis reveals a correlation between the time-course of the h-current’s contribution to the inward current prior to spiking and features of the FDG. By limiting our alterations to the model neurons *only* to the *Ih* maximum conductance, we can confidently conclude that the activity of the h-channel is responsible for these changes. In turn, we can directly infer a relationship between the h-channel’s expression (as quantified by the *Ih* maximum conductance), features of the h-current’s activity prior to spiking, and the presence (or lack thereof) of key features of the FDG of human L5 cortical pyramidal neurons.

### Comparison to rodent models identifies effect of h-current kinetics on FDG profile

The above analysis uncovers a relationship between the distinctive contribution of the h-current to neuronal dynamics prior to spiking and the features hallmarking the FDG of human L5 cortical pyramidal neurons. We obtained this relationship by directly comparing the human cell model’s output and analogous experimental outputs expressing complex dynamics, combined with novel analyses of the model’s dynamics. Given this and the known differences between rodent and human cortical cells, we wondered whether this relationship would be preserved for rodent cortical pyramical cells. While a biophysically-detailed model of a rodent L5 pyramidal neuron is available Hay et al. (2011), an experimental quantification of these neuron’s FDG was not. We thus performed new *in vitro* explorations.

These experiments reveal clear differences between the FDGs of rodent and human L5 cortical pyramidal neurons: notably, the primary peak in rodent L5 pyramidal neurons occurs at approximately 8 Hz, rather than between 2-6 Hz in human neurons. Additionally, there are no significant differences between the curves before and after the application of ZD7288 in rodent neurons (Figure 6**A-B**). These inter-species differences are emphasized in Figure 6**E**, including significant differences between the settings before the application of ZD7288 around the low-frequency peak of the human neurons.

**Figure 6:**
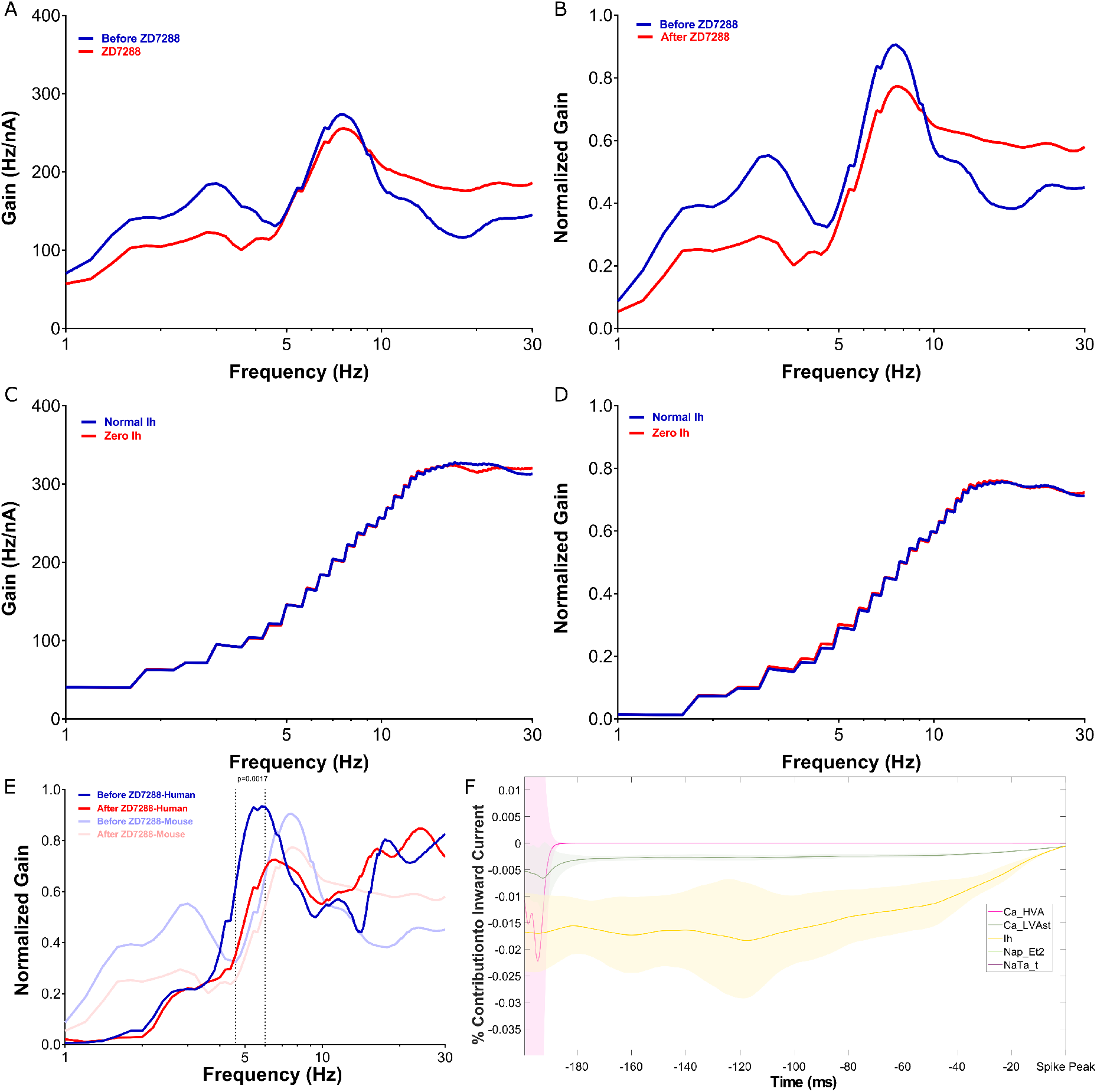
FDG properties are distinct in rodent L5 cortical pyramidal neurons, both *in vitro* and *in silico*, corresponding with a distinct h-channel contribution. **A-B** Non-normalized (**A**) and normalized (**B**) FDGs averaged from *n* = 6 rodent L5 cortical pyramidal cells firing approximately in the theta frequency range. **C-D** Non-normalized (**C**) and normalized (**D**) FDG analysis of the rodent L5 model of Hay et al. (2011) captures key qualitative properties displayed experimentally. Notably, there is minimal change in the FDG when the h-current is blocked, a stark contrast from the human setting both *in vitro* and *in silico*. The qualitative “jaggedness” of this curve relative to those from the human model (Figure 1**C-D**) are due to computational limitations described in the Methods, but do not affect the primary conclusions drawn from this analysis. **E** Comparison between the normalized FDGs of human and rodent L5 cortical pyramidal neurons highlights their differences; these are statistically significant before treatment with ZD7288 with p=0.0017 between 5-5.4 Hz (2-way ANOVA-Bonferroni’s multiple comparisons test). **F** STA analysis of the inward currents for the default model of Hay et al. (2011) (magnified to emphasize the contribution of the h-current). In comparison to the human model, the contribution is both much smaller in magnitude and notably more monotonic.

The rodent model of Hay et al. (2011) (Figure 6**C-D**) preserves some key features of the experimentally quantified FDG, including that these neurons do not exhibit a low frequency peak and exhibit minimal differences with and without h-current activity. While this correspondence is less rigorous than in the human setting, this is to be expected considering the modeling process of Hay et al. (2011) was distinct from that of Rich et al. (2021), including a reduced emphasis on capturing h-current driven subthreshold activity. Nonetheless, this correspondence between the *in silico* and *in vitro* settings highlights that we can reasonably justify using this model in similar analyses as performed on the human model to assess the relationship between ion channel contributions and FDG properties.

We note that the minimal effects of h-channel blockade on rodent neurons, either using ZD-7288 *in vitro* or by making the *Ih* maximum conductance zero *in silico*, is supported by existing literature. In a comparative study of rodent and human Layer 2/3 (L2/3) cortical pyramidal neurons, Kalmbach et al. (2018) found application of ZD-7288 had a notably diminished effect on rodent neurons relative to human neurons, including no appreciable change in the rodent neurons’ resting membrane potential and a diminished change in their input resistance. Our new experiments add to these findings by showing that ZD-7288 has a non-significant effect on the FDG of rodent neurons, while this effect is significant in human neurons.

With the appropriateness of the Hay et al. (2011) model thus justified, we applied our STA/Currentscape analysis to analyze the contribution of inward currents, particularly the h-current, to neuronal dynamics in the moments prior to spiking. This reveals notable differences in the Currentscape STAs of the rodent model compared to the human. First, the primary contributors to neuronal dynamics are the *Na_Ta_t* and *Ca_LVAst* currents (as opposed to the *Na_Ta_t* and h-currents in the human model), both of which have largely monotonic dynamics upon qualitative inspection in the 200 ms prior to spiking. Second, the h-current’s contribution is notably smaller, meriting the magnification presented in Figure 6**F**. Finally, the contributions of the *Nap_Et2* and *Ca_HVA* currents are smaller by approximately another order of magnitude and largely negligible.

The dynamics of the h-current’s contribution appear to be more mononotonic than in the human model upon qualitative inspection. We confirmed this using the same quantification as performed on the human model, finding that 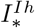 decreases for 84.56% of the STA time-series, more often than the 62.50% in the human setting.

Implicit in the overall hypothesis of this work, that the distinguishing features of the h-current’s contribution to neuronal activity are necessary for the distinct features of the FDG of human L5 pyramidal neurons, is the inverse: neurons with different h-current dynamics should not exhibit these FDG properties. Our analysis of the Hay et al. (2011) model and rodent *in vitro* results showcases exactly this phenomenon: rodent L5 pyramidal neurons lack low-frequency FDG peaks both *in vitro* and *in silico*, which corresponds with a diminished and more monotonic contribution of the h-current to neuronal dynamics in the 200 ms prior to spiking. In essence, we here have adapted the strategy of “proof by contradiction” to support our main hypothesis via a convincing example in which the inverse of our argument holds.

## Discussion

In this study, we injected a noisy input current into a biophysically-detailed, multi-compartment model of a human L5 cortical pyramidal neuron, extracting precise characterizations of ionic current contributions to spiking activity that are not attainable with analogous *in vitro* experiments. This yields three major conclusions regarding the relationship between the h-current and the frequency preference of these neurons: 1) Disruption of h-current activity alone dissipates the low frequency peak in the FDG that hallmarks these neurons’ spiking frequency preference; 2) The h-current exhibits distinctive activity in the 200 ms leading up to spiking, a previously undescribed phenomenon providing a viable explanation for the neuron’s frequency preference at this time scale and its dependence on h-current activity; and 3) Differences in this frequency preference between human and rodent L5 pyramidal neurons correspond not only with distinct h-current kinetics between species (Rich et al., 2021; Kole et al., 2006), but also the h-current’s activity in physiologically relevant settings. This supports the recent hypothesis (Moradi Chameh et al., 2021), driven by *in vitro* experiments, that the h-channel’s contribution is necessary for the frequency-preference of human L5 cortical pyramidal cells and their “leading” role in human cortical theta oscillations (Florez et al., 2015; McGinn and Valiante, 2014).

The *in silico* setting is not only useful, but in fact necessary, to derive these conclusions, as it is not possible experimentally to simultaneously record a neuron’s membrane potential and multiple ionic currents in response to a complex input. This is particularly relevant for studying the h-current, as during *in vitro* current-clamp experiments, properties of this current are most commonly inferred from the “sag voltage” (Moradi Chameh et al., 2021; Kalmbach et al., 2018; Zemankovics et al., 2010; Guet-McCreight et al., 2021) rather than directly quantified. We fully exploited the opportunities presented by the *in silico* setting by combining a contemporary visualization tool, Currentscape (Alonso and Marder, 2019), with ubiquitous spike-triggered average analysis (Schwartz et al., 2006) to identify the average contribution of each ionic current, including the h-current, in the moments prior to spiking in response to a noisy stimulus. Our computational models allowed us to manipulate the ionic current contributions in a precise fashion, unlike what would be possible in *in vitro* settings with pharmacological blockade (which also typically have secondary effects). This facilitated our connection between the enhanced non-monotonicity of the h-current’s dynamics when the *Ih* maximum conductance was doubled, and the mitigation of this dynamic when this conductance was halved, with corresponding changes in the FDG to further support our conclusions. As these were the only changes implemented in the model neuron, it is apparent that these changes to the h-current directly drive the observed changes in neuronal activity. Similar Currentscape/STA approaches have been used with a computational model of a hippocampal interneuron to show the underlying biophysical currents contributing to theta frequency spiking resonance (Sun et al., 2022).

The correspondence between the *in vitro* FDG (Moradi Chameh et al., 2021) and that of the model neuron of Rich et al. (2021) is vital and non-trivial. Given inherent limitations in the study of human cortical tissue, the model of Rich et al. (2021) was generated primarily by matching subthreshold voltage activity from a single neuron in a range with pronounced h-current activity, including adjusting the h-current’s kinetics. Spiking dynamics were constrained only by requiring the neuron to fire repetitively in an approximate frequency range exhibited by separate human L5 neurons in response to tonic inputs. The model’s close approximation of complex spiking dynamics observed *in vitro*, but not constraining model development, is additional support that the h-current plays a pivotal role in this activity.

We note that our analysis of the *in silico* model is limited to the soma to correspond to the *in vitro* experiments. The frequency preference of dendrites is a rich topic for future research given both the non-uniform distribution of the h-channel (Rich et al., 2021; Hay et al., 2011) and unique morphological properties of human neurons (Beaulieu-Laroche et al., 2018). We also acknowledge the limitations inherent in all computational modeling studies: the model neurons studied here are necessarily abstractions of the biophysical reality that cannot capture the entirety of a neuron’s intricate dynamics (Almog and Korngreen, 2014). Nonetheless, the focused conclusions of this study on the relationship between the h-current and spiking frequency preference are well justified given the specific correspondences between the *in vitro* and *in silico* settings in h-current activity and FDG. Given neuronal degeneracy (Edelman and Gally, 2001), it is possible that combinations of other ionic currents might affect the FDG properties related to h-current activity in this study; however, that does not affect the primary conclusion of this work, that that the h-current is a necessary contributor to these FDG properties for this well-developed model of a human cortical L5 cell. Developing and exploring additional models and model populations is outside the focused scope of this paper.

We have shown here that h-channels are essential in determining the spiking frequency preference of human L5 pyramidal cells, which in turn implies that they can be key controllers of oscillatory activity produced by neuronal circuits. As more human data has become available, computational models of human cortical circuits have been developed and used to show how reduced inhibition in depression affects stimulus processing (Yao et al., 2022), how reduced neuronal heterogeneity can impair seizure resilience (Rich et al., 2022), and that age-dependent sag current increases affect resting state activities (Guet-McCreight et al., 2021). While circuit models are needed to link to system level experimental data, there are many more experimental unknowns that need to be considered in developing such models. Indeed, exploration goals using model neuronal circuits often differ from those of studies using detailed cellular models of individual neurons. This work is an example of the latter, focusing on whether and how the specifics of particular channel types might affect complex cellular output (i.e., spiking frequency preference) that influence circuit output (i.e., rhythms). The detail needed in any model naturally varies with the question, and there is no clear consensus of the extent of experimental detail needed as we consider the multiscale nature of the brain (D’Angelo and Jirsa, 2022). Given these challenges, clarity regarding model generation and the goals of computational studies are required to accelerate our understanding of the brain (Eriksson et al., 2022) and usher interdisciplinary neuroscientists towards jointly tackling the challenges of neurodegenerative disease at cell and circuit levels (Farrell et al., 2019; Gallo et al., 2020; Rich et al., 2022).

## Acknowledgements

Funding sources: the National Sciences and Engineering Research Council of Canada (RGPIN-2016-06182 to F.K.S., RGPIN-2015-05936 to T.A.V.), the Krembil Foundation, the University of Toronto Department of Physiology, and the Savoy Foundation (fellowships/awards to S.R.).

## Notes

**Conflicts of Interest**: The authors declare no competing financial interests.

### Competing Interest Statement

The authors have declared no competing interest.

